# Environmental and ecological signals predict periods of nutritional stress for Eastern Australian flying fox populations

**DOI:** 10.1101/2023.12.01.569640

**Authors:** John Lagergren, Manuel Ruiz-Aravena, Daniel J. Becker, Wyatt Madden, Lib Ruytenberg, Andrew Hoegh, Barbara Han, Alison J. Peel, Peggy Eby, Daniel Jacobson, Raina K. Plowright

## Abstract

Food availability determines where animals use space across a landscape and therefore affects the risk of encounters that lead to zoonotic spillover. This relationship is evident in Australian flying foxes (*Pteropus* spp; fruit bats), where acute food shortages precede clusters of Hendra virus spillovers. Using explainable artificial intelligence, we predicted months of food shortages from climatological and ecological covariates (1996-2022) in eastern Australia. Overall accuracy in predicting months of low food availability on a test set from 2018 up to 2022 reached 93.33% and 92.59% based on climatological and bat-level features, respectively. Seasonality and Oceanic El Niño Index were the most important environmental features, while the number of bats in rescue centers and their body weights were the most important bat-level features. These models support predictive signals up to nine months in advance, facilitating action to mitigate spillover risk.

## Introduction

Food availability drives animal distribution across landscapes with cascading effects on their demography, human-wildlife conflict, and zoonotic spillover. Human-induced pressures, such as land conversion and climate change, alter resource quality, abundance, and reliability, often resulting in food scarcity (Birnie-Gauvin *et al*. 2017). Predictive models that forecast resource scarcity experienced by wildlife may be leveraged to predict and mitigate the consequences of changing food availability, such as zoonotic spillover.

As nectarivores and frugivores, pteropodid bats disperse pollen and seeds (Aziz *et al*. 2021), helping to regenerate and maintain plant diversity. In recent decades, pteropodid numbers have declined, with many species now listed in categories of conservation concern. Many pteropodid species are nomadic and track ephemeral but high-yield food across large distances (Giles *et al*. 2016), making them vulnerable to food scarcity or habitat change (Eby 1991). Additionally, land clearance has removed most of the winter-flowering resources used by Australian *Pteropus* species (flying foxes), forcing bats to rely on alternative but nutritionally suboptimal resources (Eby 1998; Eby *et al*. 1999; Nelson *et al*. 2000).

Food shortages affect bat physiology and behavior and, consequently, infection dynamics (Parry-Jones *et al*. 2016; Eby *et al*. 2023). Australian flying foxes buffer native food scarcity by foraging on alternative resources in human-modified areas (Markus and Hall 2004; Williams *et al*. 2006; Meade *et al*. 2021; Eby *et al*. 2023). This increases contact with horses, facilitating spillover of Hendra virus, a zoonotic pathogen for which flying foxes (*P. alecto* in particular) are reservoirs (Plowright *et al*. 2015). Seasonal patterns induce scarcity of food in winters; however, periods of acute food shortages are also linked to El Niño/La Niña cycles (Eby *et al*. 2023). These ecological shifts have resulted in spatiotemporal clusters of spillovers after food shortages that are likely driven by increased contact with horses and increased viral shedding from bats (Becker *et al*. 2022; Eby *et al*. 2023). Thus, identifying early indicators of nutritional stress could aid *Pteropus* conservation and help preempt virus spillovers.

We applied machine learning and explainable artificial intelligence (X-AI) methods to identify reliable environmental and bat-level indicators of monthly food shortages for Australian *Pteropus* bats. We compiled data on months of nectar shortages in subtropical Australia over the past 20 years. We used gradient boosting decision trees, GBDT (Friedman 2001; Ke *et al*. 2017), and Shapley additive explanations, SHAP (Lundberg and Su-in 2017), to assess variable importance for classifying months as food shortages, and then used the identified variables in reduced models to predict food availability. Our results provide early signals of food shortages and may inform mitigation of bat-human conflict, conservation practices, and reduce virus spillover risks.

## Methods

### Data collection

We used recorded monthly food shortages based on nectar availability from bat diet plants like eucalyptus (Eby *et al*. 2022). The records and practices of commercial apiarists were used to track nectar productivity in bat diet plants as a proxy for bat food availability. Food shortages were defined as periods when nectar production was insufficient to prevent starvation in commercial beehives without supplemental feeding.

We focused on northeast New South Wales, Australia. The area was defined by the approximate origin area of flying foxes treated by the Northern Rivers Branch of the Wildlife Information Rescue and Education Service from January 2006 to February 2020. These records included species, age, sex, physical impairment, forearm length (millimeters), and mass (grams). While records for three flying fox species were available (*Pteropus scapulatus, P. poliocephalus*, and *P. alecto*), we excluded *P. scapulatus* due to its distinct seasonal behavior and presence in the area. We considered monthly counts of bats collected for rehabilitation (RN) and counts from 3-month and 6-month lags and cumulative totals, yielding five variables. Shorter time increments demonstrated high correlation with absolute Pearson correlation >0.95 between adjacent months and thus were excluded.

We derived monthly proxies of population-level health using 3,236 intake records. First, we used raw data, including mean weight of adult bats per species and sex. Minimum and maximum weight were highly correlated with mean weight, so the latter was kept for analysis. Given health issues of rescued bats, we determined individual body mass scores based on weight and size compared to weight-forearm length relationships in a distinct dataset of 2,205 healthy wild bats caught between 2017 and 2020 (SI Figures 3-6). This metric was obtained for 796 adult bats (494 *P. alecto* and 302 *P. poliocephalus*) with records of weight pre-treatment. We included season as a lifecycle proxy (0: summer [Dec-Feb.], 1: autumn [Mar-May], 2: winter [Jun-Aug.], and 3: spring [Sep-Nov.]). The monthly bat-level dataset comprised five rehabilitation intake numbers with cumulative or lagged values, four average mass values (two species and two sexes), mean and proportion of bats below body mass score, and season, resulting in 12 bat-level variables spanning June 2006 to February 2020 (165 months, 16 food shortages).

We compiled monthly environmental data, including three global indices (Oceanic Niño Index [ONI], Southern Oscillation Index [SOI], and Southern Annular Mode [SAM]) and five local weather variables (average maximum and minimum daily temperature [TMAX, TMIN], average daily precipitation [PPT], and heat days [HD]; SI Table 1). Temperature and precipitation were obtained from NOAA gridded climate data using the ncdf4 R package (version 1.21, Pierce 2023). Heat days were derived by summing the number of days where maximum temperature exceeded the adjusted monthly average by 5 °C, a threshold relevant for eucalyptus species growth (Araujo et al. 2003). These microclimatic variables were extracted for a forest pixel representing native bat foraging habitats in the study area, and were averaged (TMIN, TMAX) or totaled (PPT, HD) over 30-, 91-, 183-, 273-, and 365-day lags. Global indices were lagged by 3, 6, 9, and 12 months. Finally, season was included as a proxy for environmental seasonality, resulting in 36 monthly environmental variables spanning January 1996 to December 2022 (324 months, 23 food shortages).

Multiple bat-level timepoints contained missing data, particularly in early years due to recording changes. Months with >50% missing values were removed. Otherwise, missing values (0.8% of environmental and 23.7% of bat-level datasets) were imputed via Multivariate Imputation by Chained Equations (MICE) using the scikit-learn (version 1.2.2) Python package (version 3.11.2).

### X-AI approach

We applied X-AI methods to classify each month in the time series as presence (1) or absence (0) of food shortage based on environmental and bat-level features. We used gradient boosting decision trees (GBDTs) due to their ability to discover high-order interactions among input features for fast and accurate prediction together with SHAP to rank features by importance according to their contributions to each model output. Both methods were implemented in Python using the lightgbm (version 3.3.5) and shap (version 0.42.1) packages.

We used time series cross-validation to ensure future timepoints were not used to train models evaluated on past timepoints. Cross-validation folds consisted of two consecutive years for validation and all preceding timepoints for training. The first four years were used only for training, while years 2018 and onward were reserved for final model evaluation (SI Figure 1). Cross-validation folds were used to conduct a grid search over 2,187 hyperparameter combinations to prevent overfitting (SI Table 2). We limited models to tree depth of four and trained for 100 iterations. Weighted binary log-loss was used as an objective function, where weights were chosen to address class imbalance (sparse presence of food shortages over time). Optimal hyperparameter combinations were chosen to minimize the weighted average validation error, with each validation set weighted proportionally by sample size in its training set (i.e., validation scores with more training data were weighted more highly than those with fewer training samples). Final models used the best hyperparameter combinations and were trained on all data up to 2018.

We used SHAP to assess each feature’s contribution to predicting food shortages. Using GBDT as a classification model, SHAP values were represented as logits. We considered SHAP values for training and test sets. Using the SHAP values for the training set, we identified individual and pairwise feature thresholds based on Gini impurity, which informed decision rules to predict monthly food shortages.

We quantified GBDT predictive accuracy using precision-recall curves of the mean probability thresholds that maximized F-scores across validation splits. Using these optimal thresholds, we calculated classification accuracy, true positive, and true negative rates to account for class imbalance.

## Results

### Model comparison

The GBDT used probability thresholds to classify a month as food shortage of 0.3242 and 0.9869 for the environmental and bat-level datasets, respectively (Figure 1; Table 1).

**Table 1.** Monthly food shortage classification accuracy of the GBDT for the environmental and bat-level datasets. We report overall and class-level accuracies for the validation and test sets. The true positive rate (sensitivity) and true negative rate (specificity) report the proportion of correct classifications for the presence and absence of food shortages, respectively. Classification accuracy is reported at month and season level. Note that if any month in a given season was recorded as a food shortage, then the season was labeled as a food shortage.

| Environmental dataset |  |  |  |
| --- | --- | --- | --- |
|  | Overall accuracy | True positive rate | True negative rate |
| Validation (monthly) | 91.20% | 44.44% | 95.45% |
| Test (monthly) | 93.33% | 33.33% | 96.49% |
| Validation (seasonal) | 90.28% | 75% | 92.19% |
| Test (seasonal) | 90% | 100% | 89.47% |
| Bat-level dataset |  |  |  |
|  | Overall accuracy | True positive rate | True negative rate |
| Validation (monthly) | 97.92% | 75% | 100% |
| Test (monthly) | 92.59% | 100% | 91.67% |
| Validation (seasonal) | 100% | 100% | 100% |
| Test (seasonal) | 80% | 100% | 77.78% |

**Figure 1.**
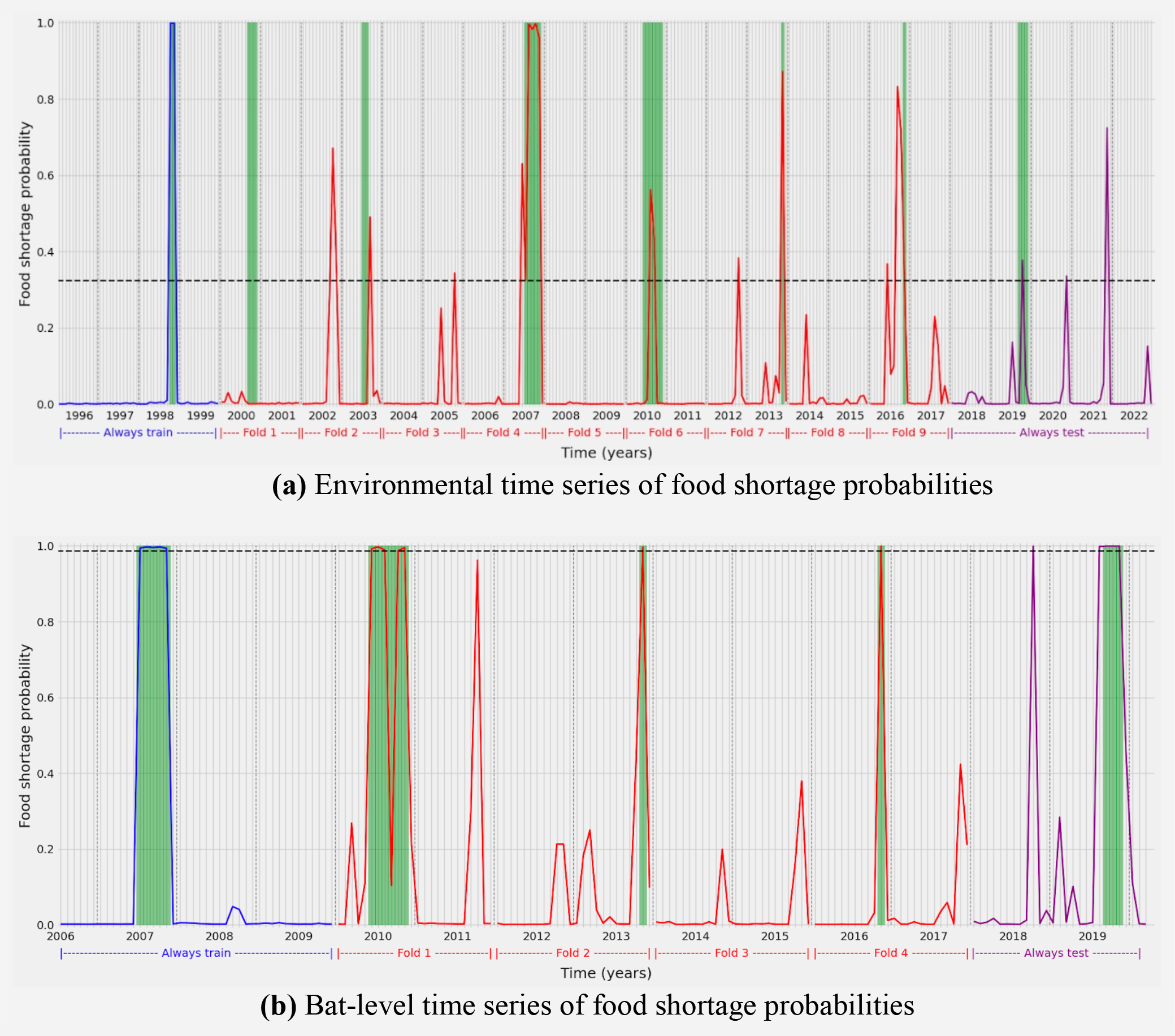
Probability estimates of monthly food shortage presence (1) and absence (0) using a GBDT classifier trained on environmental (a) and bat-level (b) datasets. The x-axis in each plot corresponds to time at monthly resolution and the y-axis corresponds to food shortage probability. The blue, red, and purple curves indicate model predictions on the training, validation, and test sets, respectively. Note that all timepoints preceding each validation fold were used for training (e.g., Fold 3 in panel (a) used data from 1996-2003 for training and 2004-2005 for validation). Green shaded regions correspond to ground truth food shortages, vertical dashed lines are used to distinguish between years, and the horizontal black dashed line is the optimal probability threshold used to classify months as food shortages.

The models performed similarly for the bat-level and environmental datasets, reporting >90% overall monthly classification accuracy for the validation and test splits (Figure 1; Table 1). The bat-level data produced a model with a three-fold higher optimal probability threshold (0.98) compared to that using environmental data (0.32). Both models accurately predicted monthly food shortage absence with >90% accuracy, however, the environmental model struggled to predict monthly food shortage presence. Aggregating monthly predictions by season resulted in an equal or better true positive rate, but with mixed results for the true negative rate. The only false negative identified by the model corresponded to the year 2000 using the environmental dataset (Figure 1).

### Drivers of food shortages

Of the 36 features in the environmental-level dataset, we identified 11 important features (with normalized mean absolute SHAP value greater than 2.5%). These included a combination of season, ONI, and others (Figure 2a). Of the 12 bat-level features, five were identified as the most predictive. These included adult male *P. alecto* mass, numbers of bats received for rehabilitation, and season (Figure 2b).

**Figure 2.**
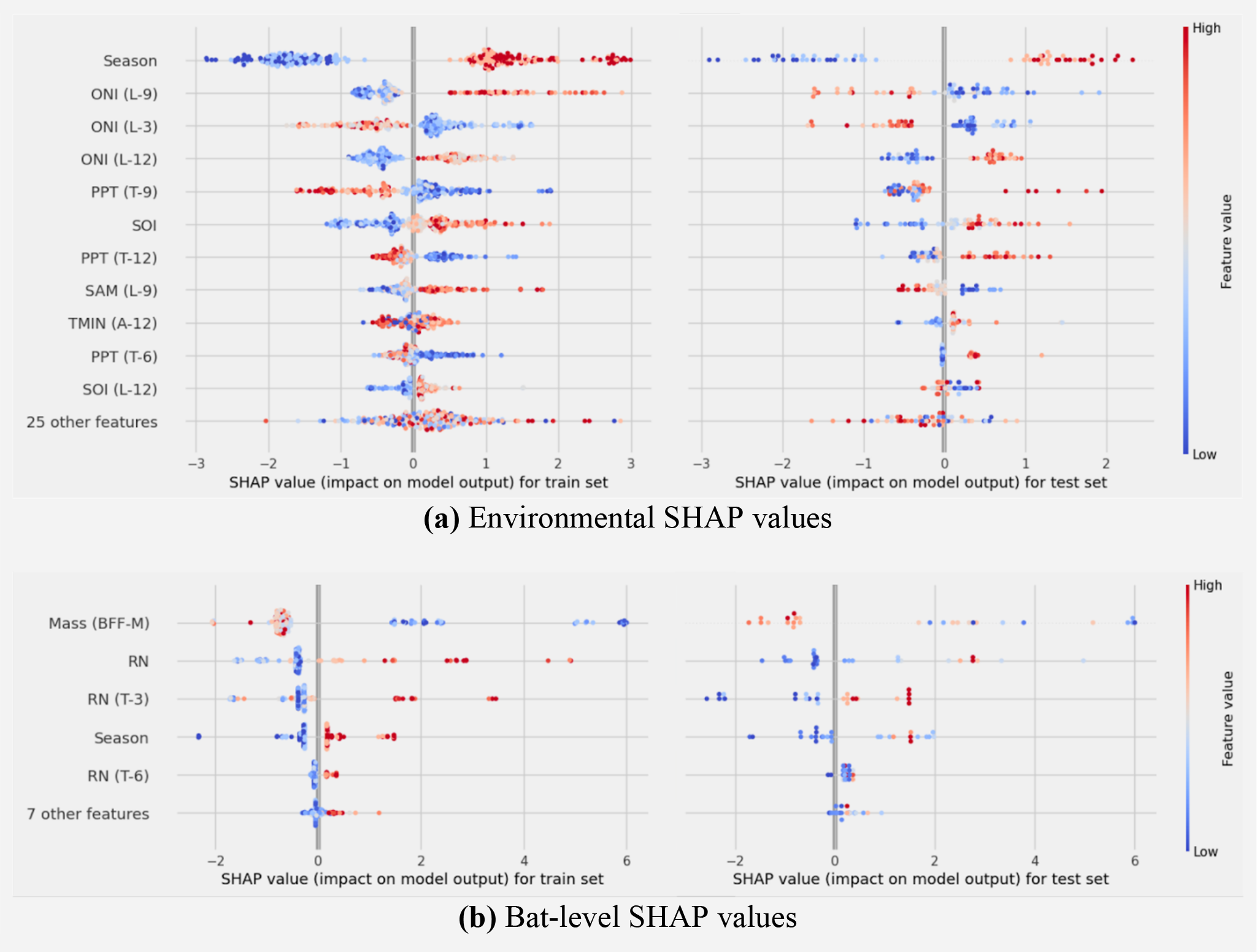
SHAP values for the environmental and bat-level training (left) and test (right) sets. The y-axis includes the most influential features identified by SHAP. The x-axis shows the SHAP value (i.e., impact on model output) in the form of logits, where negative values decrease, and positive values increase the probability of a food shortage. This plot visually represents the impact each feature has on the model’s predictions. The larger the magnitude of a SHAP value for a feature, the larger its influence on the model’s predictions. Each observation of each feature is plotted as a data point, where red and blue colors indicate large and small values of that feature, respectively. For season, low to high feature values represent summer, autumn, winter, then spring. Features where the SHAP values transition from blue to red (going left to right) are positively associated with food shortage probability, and those transitioning from red to blue are negatively associated. Glossary: L – lag, T – total, ONI – Oceanic Niño Index, PPT – precipitation, SOI – Southern Oscillation Index, SAM – Southern Annular Mode, TMIN – minimum temperature, BFF-M – black flying fox (*P. alecto*) adult males, RN – rehabilitation number.

SHAP values across the top features of both datasets revealed multiple associations between environmental and bat-level covariates with food shortages. Noticeably, top features had approximately bimodal distributions across predictions of presence and absence of food shortages (Figure 2). Seasonality presented high importance in both the environmental and bat-level datasets. Low season values (summer and autumn) produced negative SHAP values, reducing the probability of a food shortage, while high values (winter and spring) produced positive SHAP values, increasing the probability of food shortage. In the environmental dataset, longer ONI time-lags (nine and 12 months) were positively associated with food shortage probability and shorter ONI time-lags (three months) were negatively associated. In the bat-level dataset, lighter weight of adult male *P. alecto* and high intake numbers of rescued bats were predictive of monthly food shortage.

### Reduced threshold models and food shortage prediction

We focused on the top two environmental and bat-level predictors of food shortage (Figure 2) and estimated the optimal splits that separate positive from negative SHAP values (SI Figure 2). The reduced threshold models are similarly evaluated on the test set (Table 2; Figure 3).

**Table 2.** Food shortage classification accuracy of the reduced threshold models for the environmental and bat-level datasets. Each dataset includes two threshold models: one based on SHAP values for individual features and one for SHAP values for feature interactions. We report overall and class-level accuracies for the environmental and bat-level test sets. The true positive and negative rates report the proportion of correct classifications for the presence and absence of food shortages, respectively. Classification accuracy is reported at month and season level. Note that if any month in a given season was recorded as a food shortage, then the season was labeled as a food shortage.

| Environmental dataset |  |  |  |
| --- | --- | --- | --- |
|  | Overall accuracy | True positive rate | True negative rate |
| Individual (monthly) | 96.67% | 100% | 96.49% |
| Interaction (monthly) | 96.67% | 100% | 96.49% |
| Individual (season) | 95% | 100% | 94.74% |
| Interaction (season) | 95% | 100% | 94.74% |
| Bat-level dataset |  |  |  |
|  | Overall accuracy | True positive rate | True negative rate |
| Individual (monthly) | 77.78% | 100% | 75% |
| Interaction (monthly) | 85.19% | 100% | 83.33% |
| Individual (season) | 50% | 100% | 44.44% |
| Interaction (season) | 60% | 100% | 55.56% |

**Figure 3.**
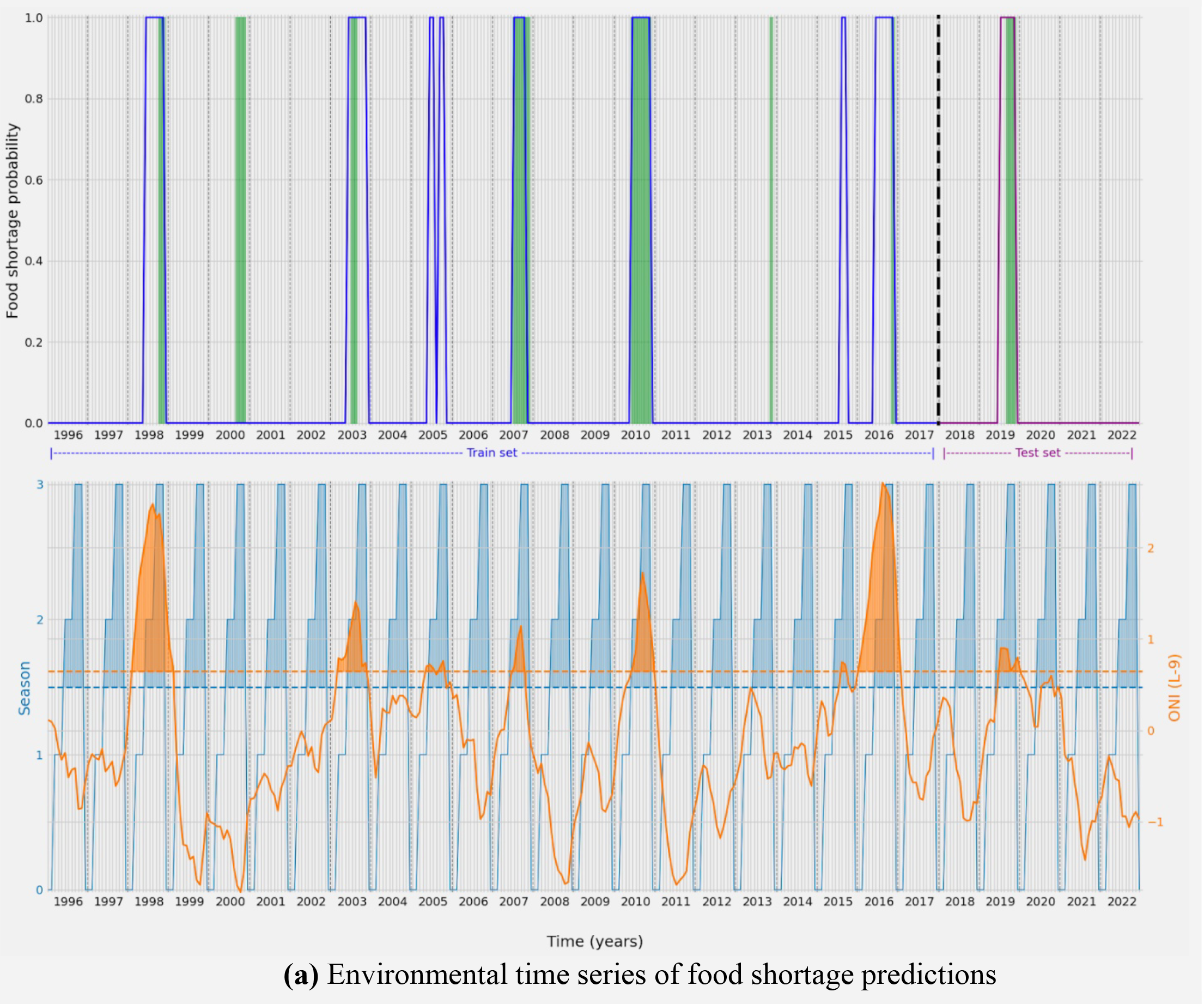
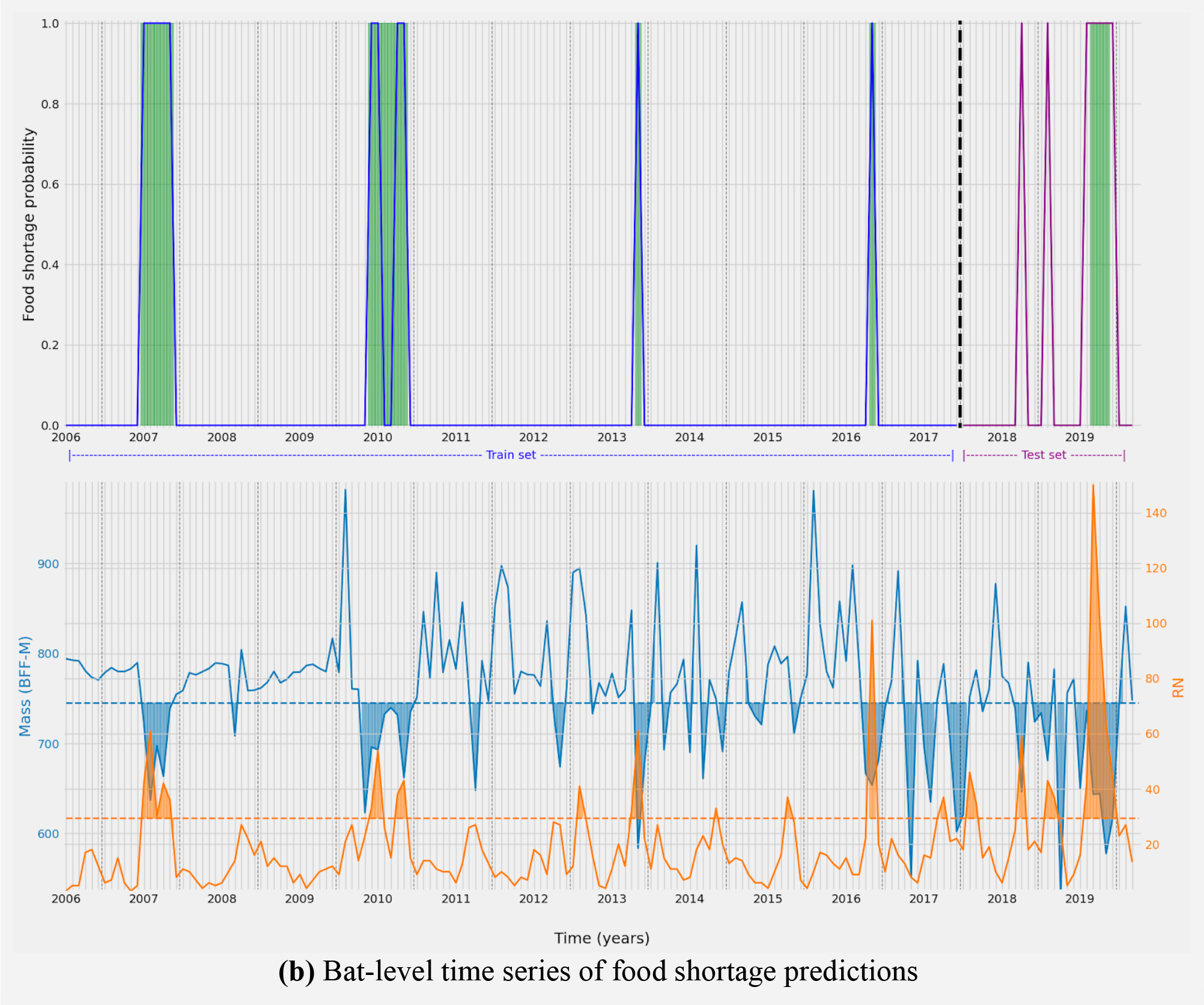
Predictions of monthly food shortage presence (1) and absence (0) using reduced threshold models for the environmental (a) and bat-level (b) datasets. The x-axis in each plot corresponds to time at monthly resolution and the y-axis corresponds to food shortage. In the top plots of (a) and (b), the blue and purple curves indicate model predictions on the training and test sets, respectively. Green shaded regions correspond to ground truth food shortages. In the bottom plots of (a) and (b), the solid blue and orange curves indicate observations of the top two features, respectively, with the optimal thresholds given by the dashed lines. Shaded regions indicate where the observed values cross their respective thresholds. Months in which both features cross their thresholds are classified as food shortages. Glossary: L – lag, ONI – Oceanic Niño Index, BFF-M – black flying fox (*P. alecto*) adult males, RN – rehabilitation number.

Seasons > 1.50 (i.e., winter and spring months) with nine-month-lagged ONI > 0.65°C positively predict food shortages. These optimal thresholds were consistent across individual and interacting feature SHAP values (SI Figure 2a). Mean body mass of adult male *P. alecto* < 745.5 g and rehabilitation number > 24 bats rescued predict food shortages for features treated individually. When considered in interaction, mean *P. alecto* male mass < 745.5 g and rehabilitation number > 29 predicted food shortage with greater accuracy (Table 2; SI Figure 2b).

Each reduced threshold model is simple, interpretable, and increases prediction accuracy of food shortages for the environmental dataset (Table 2; Figure 3). The threshold model increased all accuracy metrics for the environmental dataset, while maintaining 100% food shortage prediction accuracy but lowered overall accuracy and true negative rate for the bat-level dataset. Using SHAP values on feature interactions, rather than individual features, yielded equal or greater accuracy scores across both datasets (Table 2).

## Discussion and conclusions

Winter flying fox foraging habitats in continental Australia have dramatically reduced since the arrival of Europeans as a result of land clearing, land-use change, and wildfires. Over the past 30 years, this habitat reduction, and associated food scarcity, has resulted in ecological changes in flying fox populations (Eby *et al*. 2023). We leveraged long-term data on flying fox ecology, environment, and records from a wildlife rescue center to predict months of food scarcity; these periods were recently identified as drivers of increased virus spillover risk (Becker *et al*. 2022; Eby *et al*. 2023). By using X-AI models, we classified months of food scarcity with high accuracy, forecasting capacity, and ecological explainability. Although we did not identify mechanisms driving food availability, our models reveal quantitative predictors of these periods at different temporal scales, providing signals up to nine months in advance. Exploration of causal effects and analysis at different spatial and temporal resolutions will be valuable for further developing forecasting tools to inform management to enhance bat conservation and to limit bat virus spillover.

We identified a clear hierarchy across variables predicting periods of food scarcity, from global-scale El Niño cycles to local seasonality. These findings are consistent with the phenology of eucalyptus species in Australia while also aligning with the well-established influence of El Niño cycles on global climate that might override effects of local weather. Importantly, the nine-month-lagged ONI threshold of 0.65°C is supported by previous analyses of Hendra virus spillover in eastern Australia that have also related these events with food availability for flying foxes using a Bayesian network approach (Peel *et al*. 2017; Giles *et al*. 2018; Eby *et al*. 2023). Our results therefore support the utility of this threshold for predictive and forecasting purposes. In particular, the environmental model successfully predicted the 2012 and 2013 shortages, which were not preceded by the strong El Niño events that were otherwise predictive of food shortages (Eby *et al*. 2023). This suggests the model’s nuanced capability to anticipate non-ONI-related food shortages, indicating its potential for future iterations to spatially predict shortages across different landscape pixels based on localized environmental data.

Importantly, models using bat-level data clearly identified the impact of food shortages on bat health. Each month, the total intake of 30 or more bats received for veterinary care with male adult *P. alecto* presenting average weight less than 745.5 g positively predicted food shortages. These results suggest the total number of bats rescued and their body weights—variables that can easily be recorded by rehabilitators—could be used for real-time monitoring of bat populations to inform management actions to mitigate impacts of food shortages.

During food shortages, bats also change their behavior and search for food in peri-urban areas where susceptible horses graze, increasing spillover risk (Eby *et al*. 2023). Mitigation of food scarcity for flying foxes (e.g., strategic tree planting) could therefore protect public health by improving bat health and promoting their natural nomadic behavior.

Food availability is difficult to measure at large spatial scales, and our analyses demonstrate the value of long-term data for identifying patterns. Although our datasets were restricted to a small portion of the distributional range of flying foxes in Australia, our methods identified biologically meaningful relationships to develop accurate forecasts. These methods would benefit from expanding bat-level data used in our current framework (e.g., body weight for all individuals rescued) and incorporating data at finer resolutions (e.g., better estimates of individual body condition and local environmental variables across a broader scale). Considering the small amount of winter-flowering habitat remaining in subtropical eastern Australia (Eby and Law 2008), directed efforts to conserve and restore these habitats may mitigate Hendra virus spillovers (Eby *et al*. 2023). Further analyses that leverage high-resolution data at broader spatial and temporal scales will be crucial to mechanistically identify how food scarcity affects flying fox behavior and physiology and how restoration efforts could prevent spillover.

## Data accessibility

Datasets have been deposited in Dryad Digital Repository.

## Acknowledgements

We acknowledge the Bundjalung and Widjabul Wia-bal people, who are the Traditional Custodians of the land upon which this work was conducted.

Records in the NR WIRES database reflect the dedicated efforts of 120 volunteer wildlife rehabilitators, in particular Jodie Bawn, Julie Marsh, Kim McCully, Merryn West-Bird, and Renata Phelps. Barbara Wilkins and Leonie Byron-Jackson managed the database and thousands of members of the public reported flying foxes in need of assistance to NR WIRES.

CPC Global Temperature data was provided by the NOAA/OAR/ESRL PSL, Boulder, Colorado, USA, from their Web site at https://psl.noaa.gov/data/gridded/data.cpc.globaltemp.html. CPC Global Unified Precipitation data was provided by the NOAA/OAR/ESRL PSL, Boulder, Colorado, USA, from their Web site at https://psl.noaa.gov/data/gridded/data.cpc.globalprecip.html.

