## Supplementary material for "Environmental and ecological signals predict periods of nutritional stress for Eastern Australian flying fox populations": SI Figure

[6] WIRES

[7] Department of Mathematical Sciences, Montana State University, Bozeman, MT 59715, USA.

[8] Disease Ecology, Cary Institute of Ecosystem Studies, Millbrook, NY 12545, USA.

[9] Centre for Planetary Health and Food Security, Griffith University, Nathan, QLD 4111, Australia.

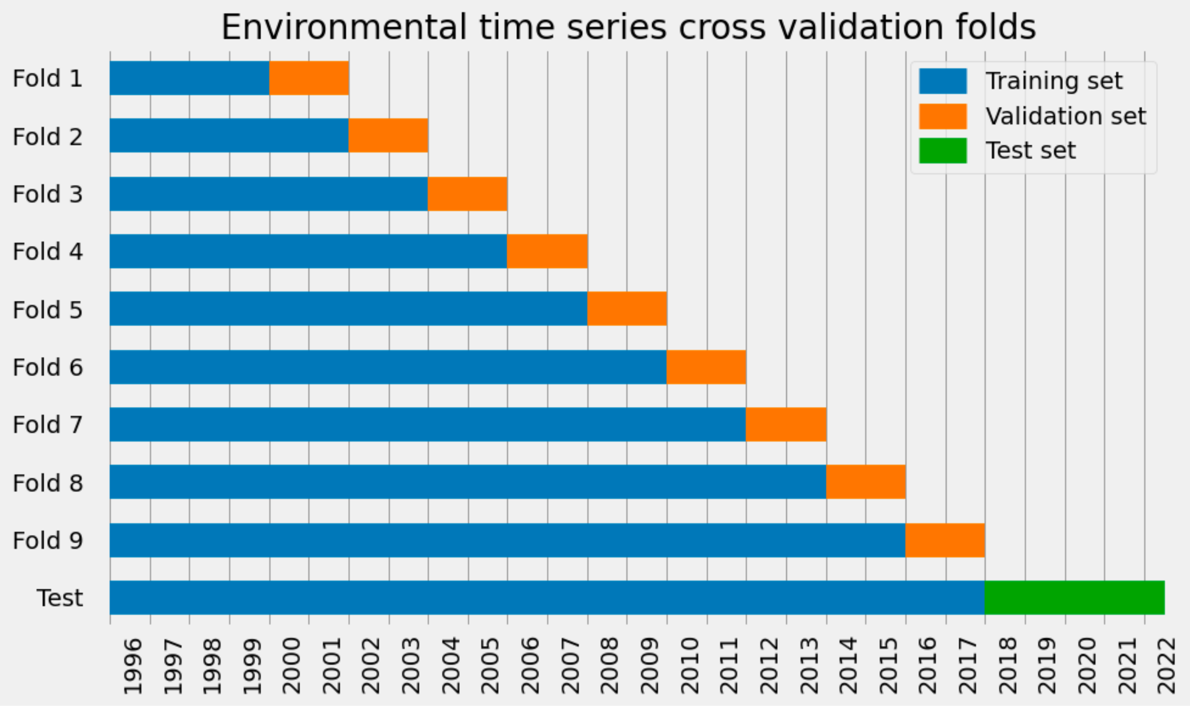


**(a)** Environmental time series cross-validation fold sets

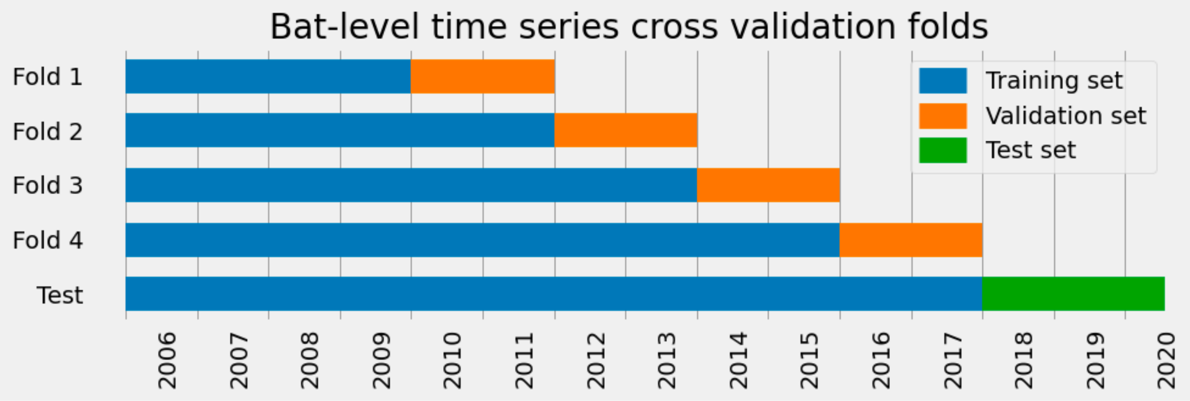


**(b)** Bat-level time series cross-validation fold sets

**Supplementary Figure 1**: Time series cross-validation folds for the environmental (a) and bat-level (b) datasets. The horizontal axis represents time in years and the vertical axis shows each cross-validation fold and test split. Training sets are shown in blue, validation sets in orange, and the final training and test split are shown in blue and green, respectively.

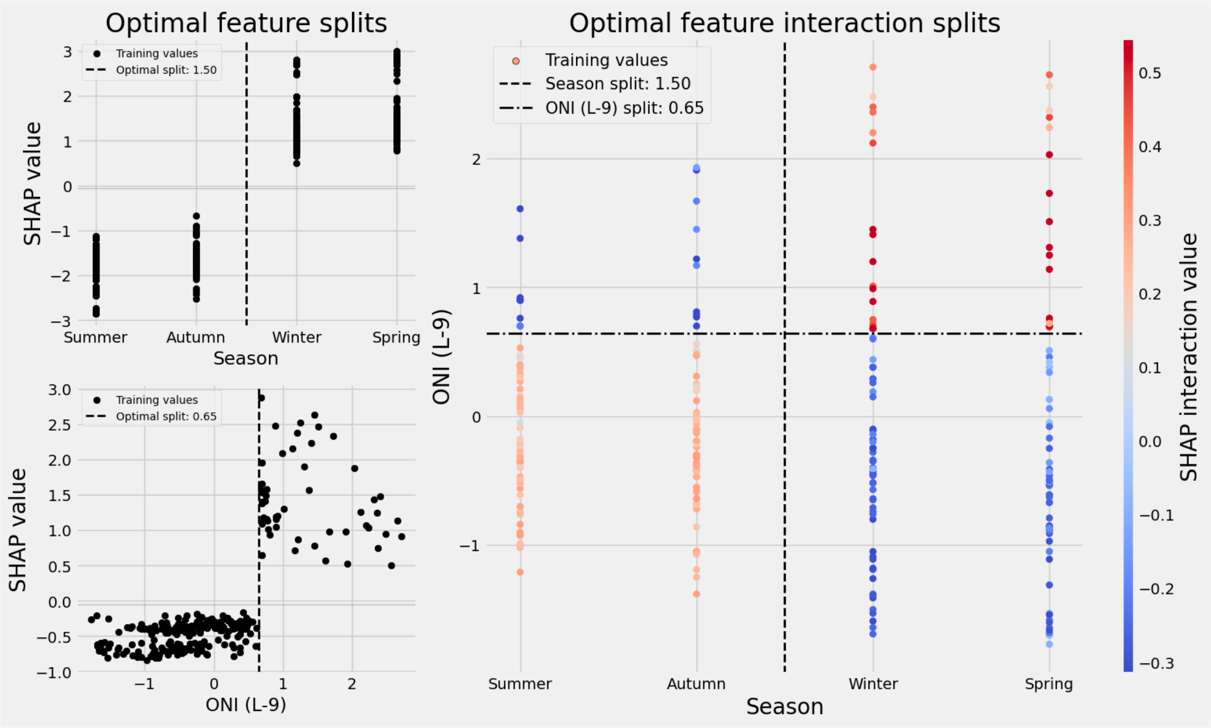


**(a)** Top environmental feature thresholds

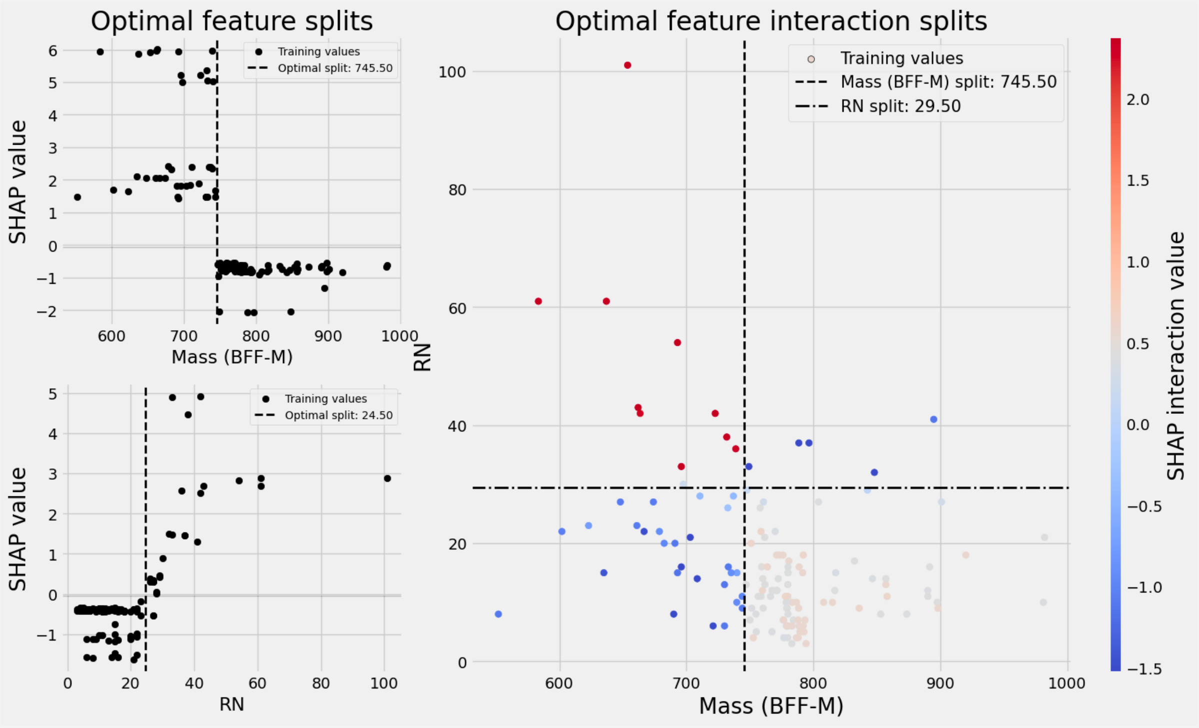


**(b)** Top bat-level feature thresholds

**Supplementary Figure 2.** SHAP values and optimal splits for individual and interacting features for environmental and bat-level datasets. Left: the y-axis corresponds to SHAP values as a function of feature observations on the x-axis. The vertical dashed line represents the optimal split that separates positive from negative SHAP values. Right: the x- and y-axes correspond to feature observations and color corresponds to the SHAP value. The vertical dashed line corresponds to the optimal split for the first top feature and the horizontal dash-dot line corresponds to the optimal split for the second top feature. The feature values for the optimal splits are given in the legend of each subplot. All plotted points belong to the training sets and the optimal splits are subsequently used for prediction on the test sets. Glossary: L – lag, ONI – Oceanic Niño Index, BFF-M – black flying fox (*P. alecto*) adult males, RN – rehabilitation number.

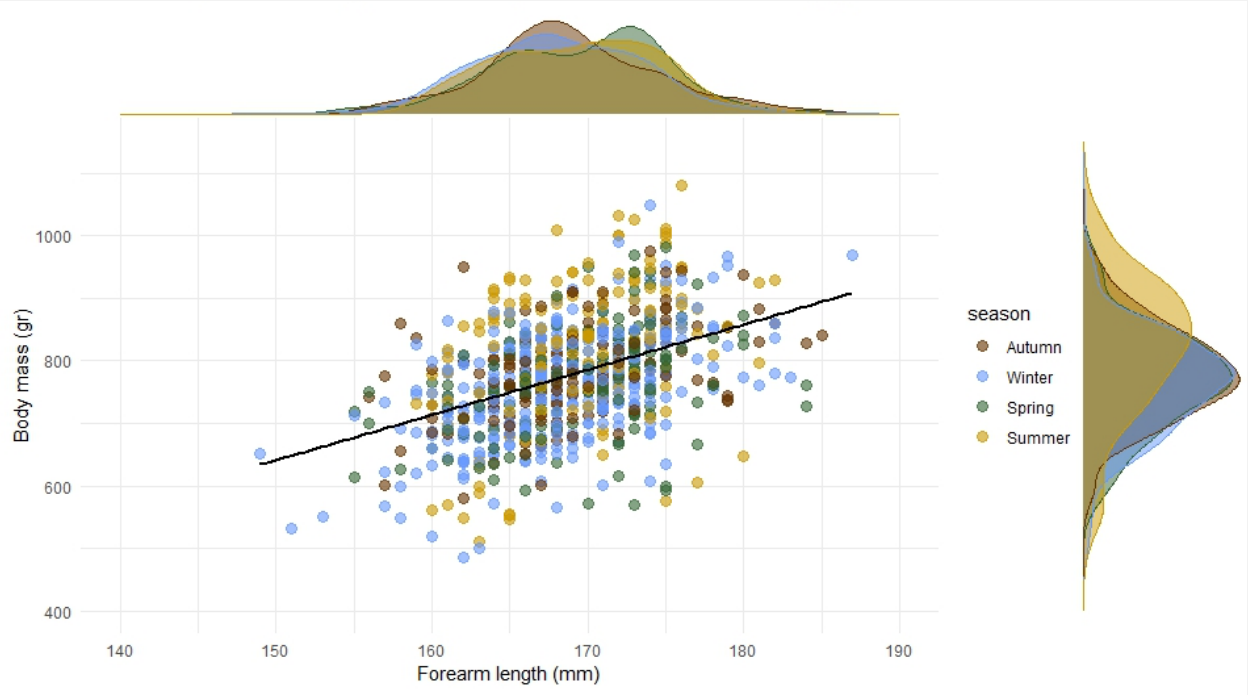


**Supplementary Figure 3:** Relationship between body mass and forearm length for adult male black flying foxes clinically healthy. Marginal distributions depict the distribution of data points across seasons.

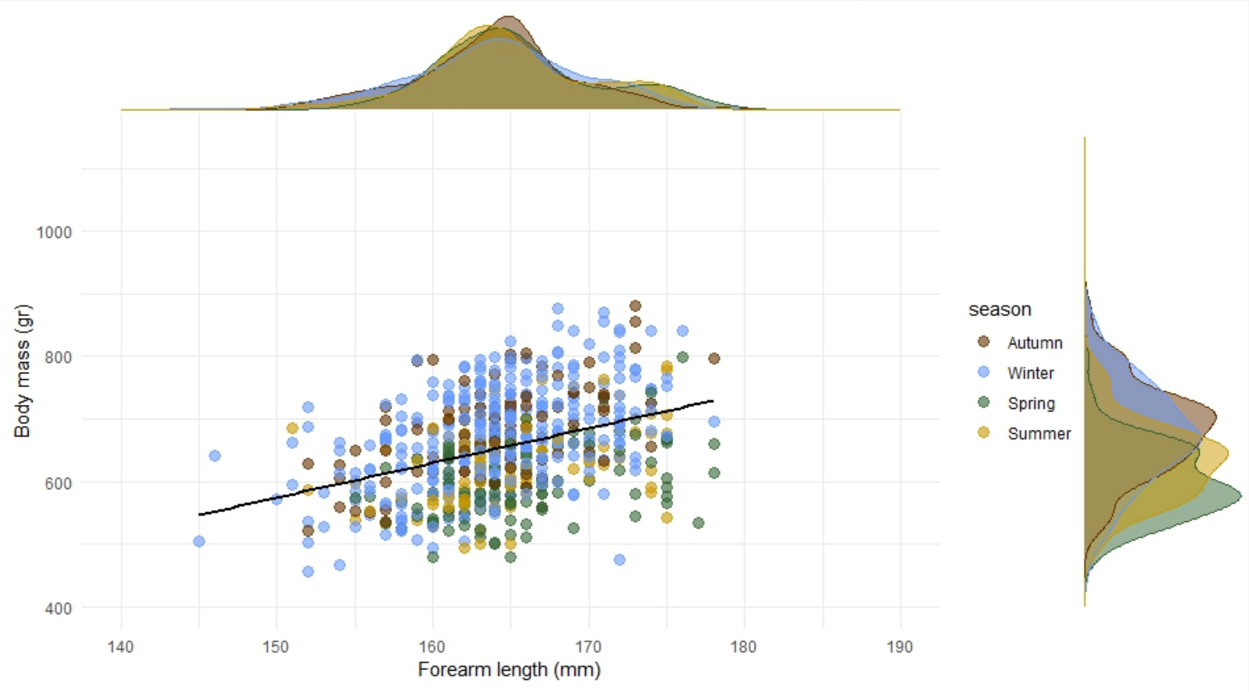


**Supplementary Figure 4:** Relationship between body mass and forearm length for adult female black flying foxes clinically healthy. Marginal distributions depict the distribution of data points across seasons.

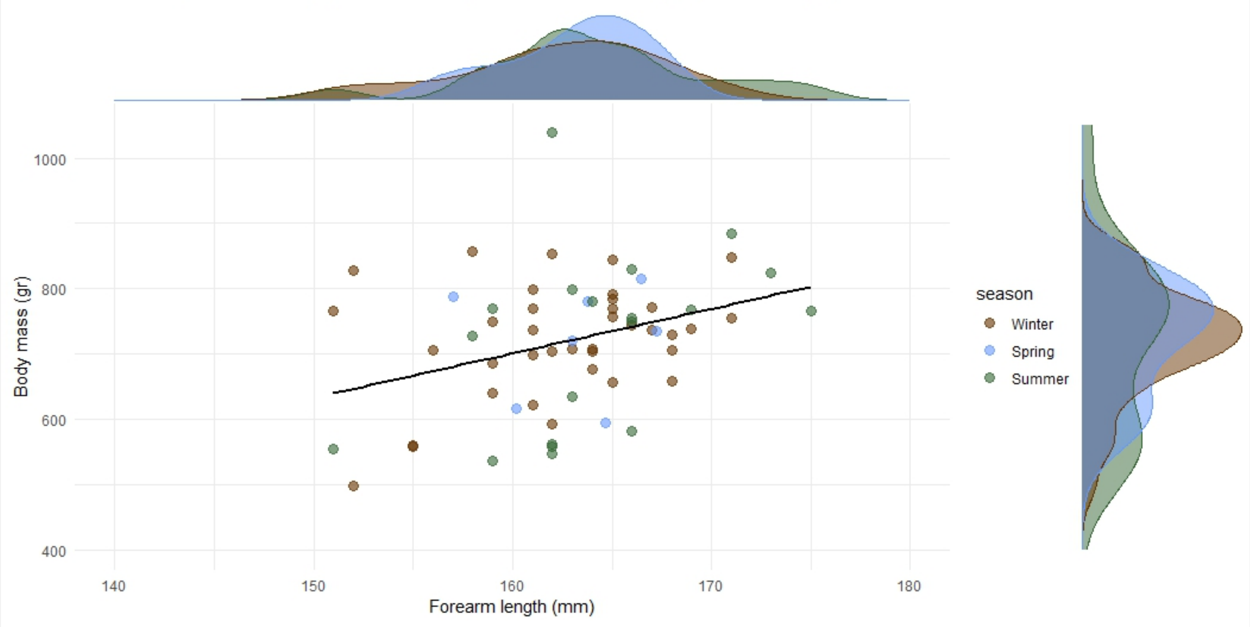


**Supplementary Figure 5:** Relationship between body mass and forearm length for adult male gray-headed flying foxes clinically healthy. Marginal distributions depict the distribution of data points across seasons.

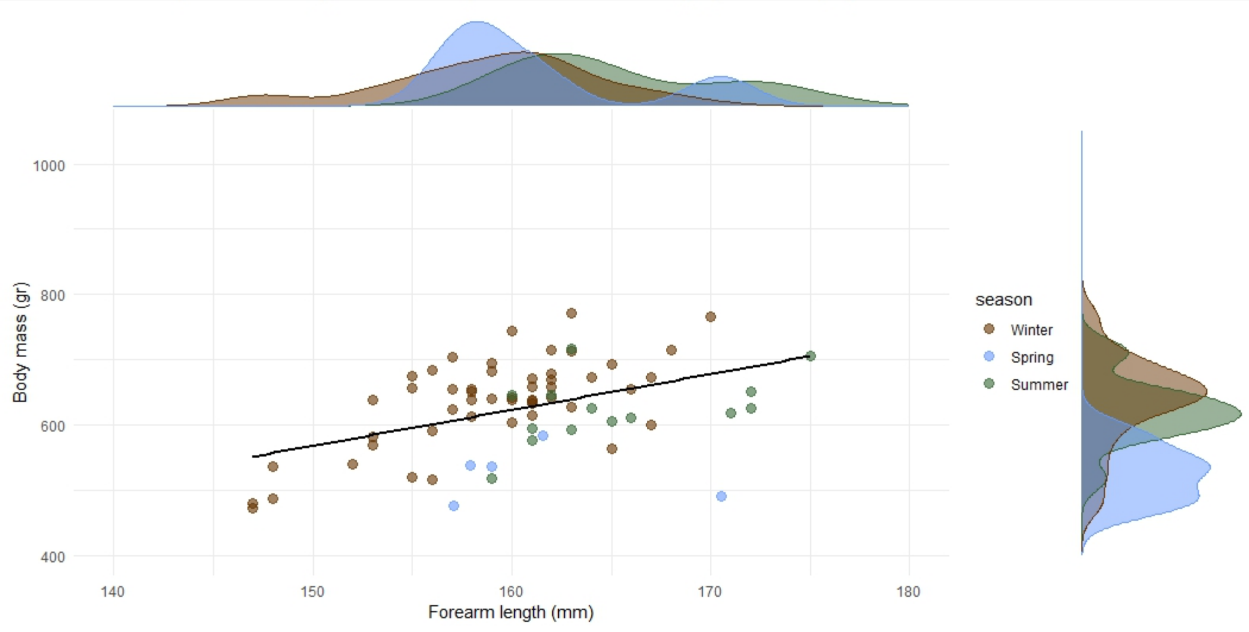


**Supplementary Figure 6:** Relationship between body mass and forearm length for adult female gray-headed flying foxes clinically healthy. Marginal distributions depict the distribution of data points across seasons.
