## Supplementary material for "Environmental and ecological signals predict periods of nutritional stress for Eastern Australian flying fox populations": SI Table

[6] WIRES

[7] Department of Mathematical Sciences, Montana State University, Bozeman, MT 59715, USA.

[8] Disease Ecology, Cary Institute of Ecosystem Studies, Millbrook, NY 12545, USA.

[9] Centre for Planetary Health and Food Security, Griffith University, Nathan, QLD 4111, Australia.

**Supplementary Table 1**: Set of environmental-level features included in the model formulations. Local variables (PPT, TMIN, TMAX, HD) were sourced from latitude -29.1722 and longitude 153.0993.

| | Units | Source | Description |
|---|---|---|---|
| SAM | - | CPC, NOAA[a] | Monthly mean Southern Annular Mode |
| ONI | °C | CPC, NOAA[b] | Monthly mean Oceanic El Niño Index |
| SOI | - | CPC, NOAA[c] | Monthly mean Southern Oscillation Index |
| PPT | mm | CPC, NOAA[d] | Total daily precipitation |
| TMIN | °C | CPC, NOAA[e] | Mean daily minimum temperature |
| TMAX | °C | CPC, NOAA[e] | Mean daily maximum temperature |
| HD | - | *Ad hoc* | Total heat days |

a Observation-based SAM index https://legacy.bas.ac.uk/met/gjma/sam.html

b https://origin.cpc.ncep.noaa.gov/products/analysis_monitoring/ensostuff/ONI_v5.php

c https://www.cpc.ncep.noaa.gov/data/indices/soi

d https://psl.noaa.gov/data/gridded/data.cpc.globalprecip.html

e https://psl.noaa.gov/data/gridded/data.cpc.globaltemp.html

**Supplementary Table 2**: GBDT hyperparameters that were tuned with time series cross-validation to prevent overfitting. Hyperparameters were chosen according to LightGBM documentation for training on small datasets.

| | Considered | Best env., bat | Description |
|---|---|---|---|
| *learning_rate* | 0.01, 0.05, 0.1 | 0.1, 0.1 | Boosting learning rate |
| *num_leaves* | 4, 8, 16 | 16, 8 | Max. leaves per tree |
| *max_bin* | 4, 8, 16 | 4, 16 | Max. feature bins per tree |
| *min_child_samples* | 1, 5, 10 | 5, 5 | Min. samples per leaf |
| *min_child_weight* | 0.001, 0.01, 0.1 | 0.01, 0.1 | Min. sum Hessian per leaf |
| *reg_alpha* | 0, 0.01, 0.1 | 0, 0 | L1 regularization penalty |
| *reg_lambda* | 0, 0.01, 0.1 | 0.01, 0 | L2 regularization penalty |
